# BrainBridge Characterizes Key Factors affecting Alzheimer’s Disease and Associated Phenotypes

**DOI:** 10.64898/2025.12.03.692255

**Authors:** Tianyu Liu, Minsheng Hao, Eric Sun, Yaroslav Markov, Le Zhang, James Zou, Hongyu Zhao

## Abstract

Single-cell RNA sequencing (scRNA-seq) has significantly advanced our understanding of Alzheimer’s disease and aging by revealing cellular heterogeneity and shifts in cell-type composition between diseased/old and healthy/young individuals. However, few existing studies utilize the rich information in single-cell transcriptomic atlases for robust patient-level modeling and biological feature selection. To address this gap, we present BrainBridge, a deep learning-based framework designed to integrate atlas-scale single-cell transcriptomic data with phenotypic information to model the biomolecular complexity of the human brain. BrainBridge functions both as a powerful predictor and an embedding model for representing sample-level expression profiles and covariates through comprehensive benchmarking. We also demonstrate its effectiveness in prioritizing key genes and cell types associated with disease progression, aging, and sex differences. We further validate our findings using external resources, including genome-wide and epigenome-wide association studies (GWAS and EWAS), spatial transcriptomics, and perturb-seq experiments. Finally, we deploy BrainBridge within an interactive, agent-powered interface that enables intuitive and user-friendly model interactions, promoting broader accessibility and application in biomedical research.

## Introduction

Aging has been shown to be associated with the initiation or progression of a number of diseases, such as Alzheimer’s disease (AD) [1, 2] and Parkinson’s disease (PD) [3]. Previous studies have shown that aging significantly affects brain function and this effect can differ between sexes [4–6]. Considering the role of the brain in nervous function and systemic health, it is important to leverage biomedical data from human brain samples to investigate the effects of aging and disease in specific brain regions. Moreover, different variations, such as the change of gene expression levels [7] or changes in cell-type proportions [8], may potentially affect the aging process and disease states, either jointly or separately. Thus, identifying the factors associated with each covariate may also reduce false discoveries in therapeutic targets [9–11].

To explore biological questions inspired by gene expression changes at cell-type resolution, researchers developed single-cell transcriptomics technologies, known as single-cell RNA sequencing (scRNA-seq) [12]. The advance of scRNA-seq allows us to profile the expression of hundreds of millions of cells [13] and further describe the heterogeneity of cell states and functions. The large number of cell profiles within and across experiments has opened the way to the discovery of new cell types [14], distinct gene programs and marker genes [15, 16], and unique patient subpopulations [17, 18]. However, most previous research has focused on isolated datasets and limited samples, and does not utilize the power of atlas-level scRNA-seq results across different studies to conduct either tissue-level or species-level analysis. As an example, in the analysis of human brain, Religious Orders Study/Memory and Aging (ROSMAP) [19] and Allen Brain Cell Atlas (ABCA) [20] are two leading scRNA-seq consortia with hundreds of samples for exploring the mechanism behind Alzheimer’s disease and the progression of chronological age, but few studies consider disease-specific or aging-specific biomarkers for better understanding disease mechanisms and treatments. Furthermore, cells sequenced from the same sample are normally recognized as having the same phenotype and metadata, such as age, sex, and disease state. However, we know very little about which genes are key factors associated with these covariates. There are studies showing that the likelihood of having AD increases with age in certain range [21, 22], while other studies have also shown differences in disease-associated genes such as *MERTK* between men and women [23]. These findings raise important questions: To what extent can we be confident that genes associated with AD are true disease-related factors, rather than confounded by aging or sex? Moreover, is it possible to identify genes that are jointly associated with both AD and aging? When investigating the correlation between molecular factors and disease phenotypes, it remains challenging to disentangle true associations from confounding influences, such as age, sex, or other biological and technical variables.

Several efforts have leveraged scRNA-seq datasets from different studies to investigate the relations between patient phenotype and single cell data, including, MultiMIL [24], MrVI [25], PILOT [26], and PaSCient [27]. MultiMIL introduces a multi-instance learning (MIL) framework [28] to predict the disease state of each cell for COVID-19 analysis. MrVI is an unsupervised deep generative model to analyze large-scale singlecell transcriptomics data with multi-sample and multi-batch settings. PILOT is also an unsupervised learning framework to refine the distance of patients with the help of scRNA-seq data. PaSCient leverages the large-scale scRNA-seq data across different cell types, tissues, and disease states to provide unified sample representations and prioritize genes and cell types for each disease state.

However, none of these methods can simultaneously model and predict an arbitrary number of covariates, as well as separate factors that can affect different phenotypes. Moreover, the phenotypes across different studies might not match precisely and may be subject to missing information or low-resolution data in the modeling process, and thus we need effective solutions to model the sample representation. Finally, all of these methods are not specifically designed for human-brain analysis and lack an accessible interface for biologists to explore. Therefore, there is a need to design an interpretable computational framework for discovering the factors associated with different phenotypes and to present the results and models in a user-friendly approach for the smooth deployment process.

Here we present BrainBridge, a novel computational framework by extending PaSCient on semi-supervised learning [29] and multi-task learning to accurately predict disease states and chronological ages, and learn the differences of factors associated with different phenotypes. By training our method based on four million cells across 345 samples from ROSMAP and ABCA, BrainBridge can model both the cell and sample representations of atlas-level brain data for AD and aging analysis, which surpasses current baseline methods in predicting and modeling biology of phenotypes. BrainBridge can also prioritize genes and cell types for AD and aging progress, and further uncover the association between selected genes and DNA methylation. Those selected signals are validated in the animal models by knocking out disease-associated or aging-associated genes [30] or observing expressed protein difference by RNAScope [31]. Finally, we deploy BrainBridge as an AI agent using large language models (LLMs) [32, 33] so that natural language can interact with biological data. Users only need to provide natural language and optional input datasets to mine the required information from BrainBridge, which could be a next-generation user-friendly tool for neuroscience research.

## Results

### Overview of BrainBridge

Our model works as a computational tool to generate sample representations and prioritize biological features associated with interested phenotypes, powered by large-scale transcriptomic data. We train BrainBridge with cohort-level transcriptomic studies in brain regions from ROSMAP and ABCA, and the target is to predict disease state, chronological age, and sex for each sample. We implement a multi-instance-task learning framework with self-consistent loss [29, 34] for robust training. The trained BrainBridge generates sample-level representations that highlight disease-enriched sample clusters and specific cell-cell interactions. Moreover, BrainBridge can also identify cells, cell types, and genes associated with AD, aging effect, as well as sex effect, which can provide a comprehensive picture to understand the linkage between different biological factors and various phenotypes. We validate BrainBridge with both external datasets and in-silico/in-vitro experiments and offer an AI-powered agent system for easier access, which further demonstrates the superiority of BrainBridge to perform as a computational tool to uncover the complexity of human brains. The corresponding landscape of BrainBridge is shown in Figures 1 (a)-(c).

**Fig. 1.**
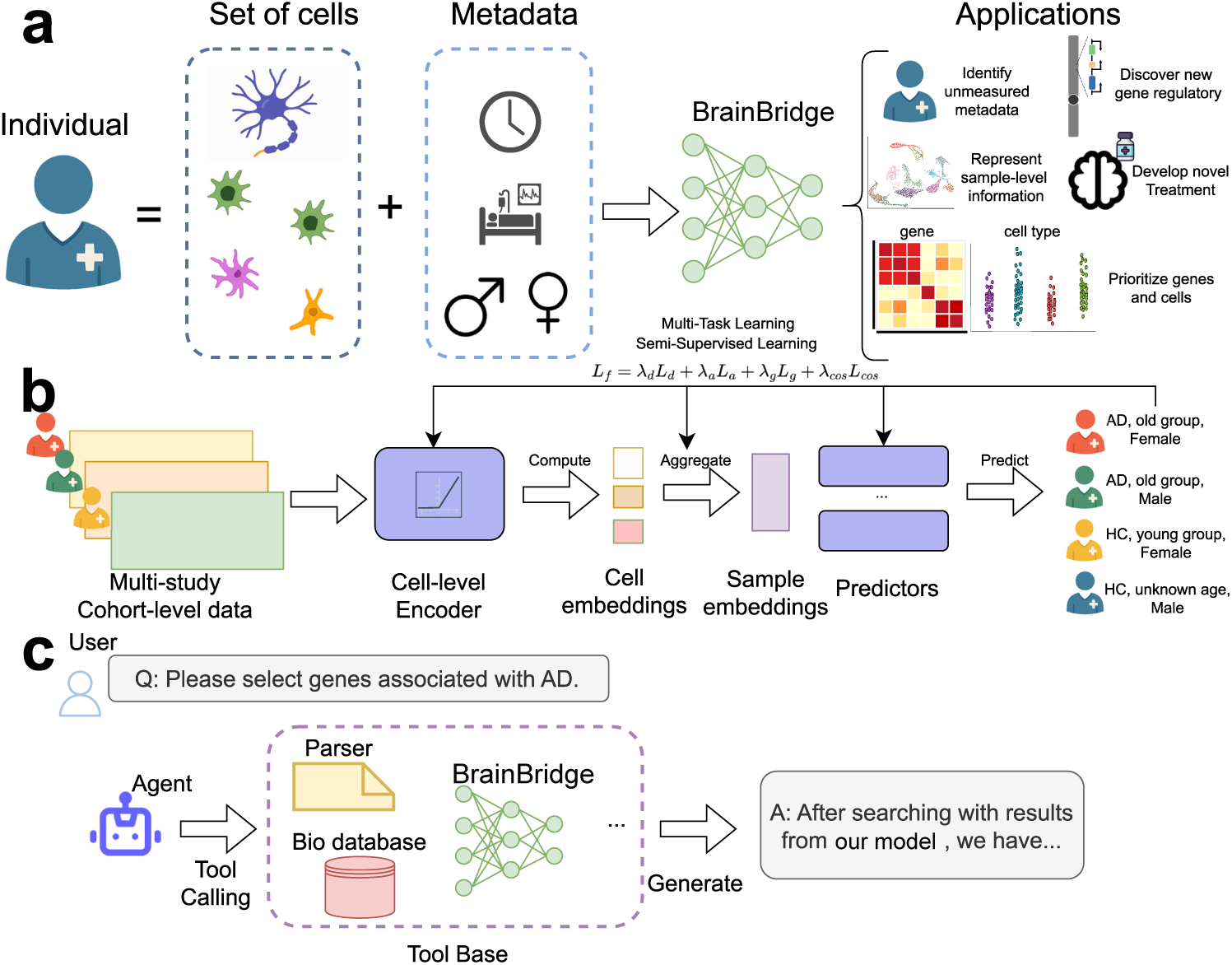
Overview of BrainBridge as a computational framework to interpret diseased samples. (a) The idea of generating patient representations from cells with phenotypes encoded in the representation. BrainBridge also supports various downstream applications for factor characterization. (b) Training BrainBridge with a multi-task learning framework. We collect multi-study single-cell atlas for training and introduce both supervised loss and semi-supervised loss for model optimization. (c) An agnetic AI system equipped with results from BrainBridge for better interaction with users.

### Benchmarking analysis demonstrates the power of BrainBridge as a predictor and embedder

We first performed a benchmarking analysis with other baseline models to demonstrate the superiority of BrainBridge in modeling different phenotypes. We first collected scRNA-seq data from ROSMAP and performed quality control. After filtering out unsatisfied cells and genes (cells: number of expressed genes smaller than 3; genes: number of expressed cells smaller than 100 [35]), we visualize the cells colored by different metadata information in Extended Data Figures 1 (a)-(d). The UMAP plots show that the ROSMAP data do not show a strong batch effect, but status signals cannot be easily distinguished visually even after applying SCimilarity [36], shown in Extended Data Figures 2 (a) and (b). The original paper of ROSMAP data leveraged MAST [37] and cell-level Wilcoxon rank-sum test to detect Differentially Expressed Genes (DEGs). To investigate the reproducibility with different pipelines, we performed DESeq2 [38] to identify disease-specific DEGs. However, only six genes were significant, recorded in Supplementary File 1. Therefore, we considered predicting disease state, sample sex, and sample age to evaluate the ability of different models as predictors. We randomly split samples from ROSMAP into training/validation/testing sets. We also inherited baselines from PaSCient including pseudobulk with transcriptomics (pseudobulk-exp), MultiMIL [24], and sample representations generated by averaging cell embeddings from SCimilarity (SCimilarity) as well as directly from PaSCient. Furthermore, we also include MrVI [25] as a new baseline method. MrVI is a computational tool developed based on scVI [39] to model sample-level covariates. Predicting disease states and sample sex are classification tasks, and thus we select Accuracy and Weighted F1 score to compare different methods. Weighted F1 score also considered label imbalance existing in the testing data. Predicting sample (chronological) age is a regression task and thus we select Mean Squared Error (MSE) and Pearson Correlation Coefficient (PCC) as metrics.

These metrics were computed with Scikit-learn [40] and Scipy [41]. We chose 10 different random seeds for conducting experiments to report variance and tuned different baseline methods with their best performances.

According to Figure 2 (a), BrainBridge outperforms other baseline methods in predicting the disease states evaluated by both Accuracy and Weighted F1 score on the testing dataset. It also surpasses the 2nd-placed method significantly, supported by the two-sided Wilcoxon Rank-sum test (p-value=0.012) [41]. Moreover, we found that baseline methods including pseudobulk-exp, Scimilarity, and MrVI consistently perform worse than BrainBridge for most seeds, demonstrating the robustness of BrainBridge as well as the limitation of generating sample representations without awareness of disease status. In Figure 2 (b), we show that BrainBridge consistently outperforms other baseline methods in predicting sample sex followed by PaSCient, which is also significantly better than the second best methods for both metrics. Therefore BrainBridge also works as a good sex classifier. Figure 2 (c) compares the results for age regression, where the predicted age from SCimilarity has a large MSE, while methods including BrainBridge, pseudobulk-exp, and MrVI generally have lower and similar MSE. By treating the averaged age from testing samples as an extra baseline method, BrainBridge also has a lower MSE compared with this baseline. Furthermore, BrainBridge has the highest averaged PCC compared with other baseline methods. We visualize the scatter plot between predicted age and observed age with BrainBridge in Extended Data Figure 3, which further supports our conclusion. By considering these factors together, BrainBridge can also work as an age clock to predict sample age from transcriptomic profiles.

**Fig. 2.**
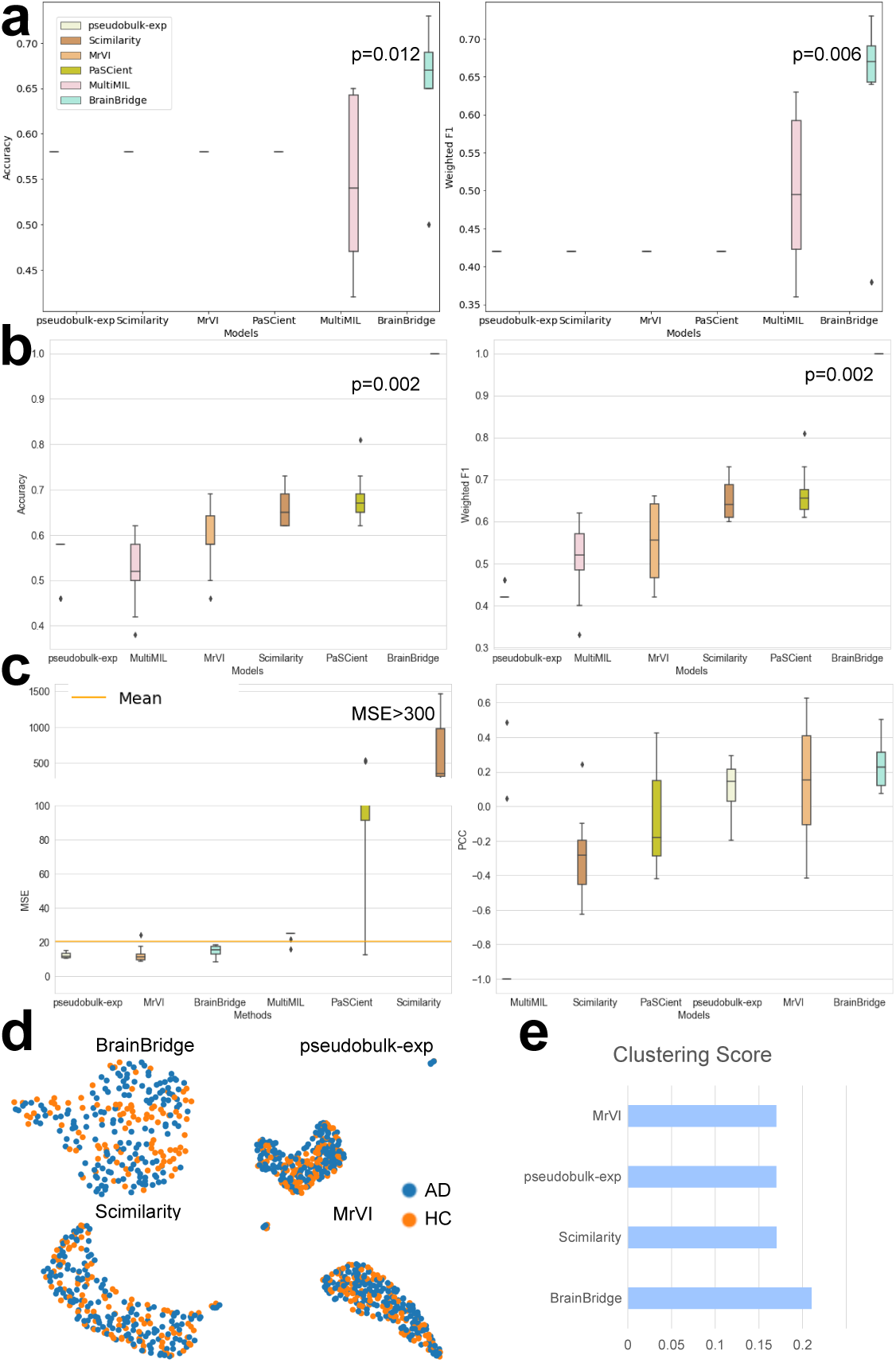
Results of benchmarking BrainBridge with other baseline models for modeling disease states, aging effect, and sex effect. (a) Comparing BrainBridge with other baseline models for predicting disease states, evaluated by Accuracy and Weighted F1 score. (b) Comparing BrainBridge with other baseline models for predicting sample sex, evaluated by Accuracy and Weighted F1 score. (c) Comparing BrainBridge with other baseline models for predicting sample age, evaluated by Mean Squared Error (MSE) and Pearson Correlation Coefficient (PCC). (d) UMAP visualization of patient embeddings generated by BrainBridge and other baselines. Each patient is colored by disease states. (e) Comparing BrainBridge with other baseline models for clustering performance. Higher scores mean better model performances for all metrics, except MSE.

We also evaluated the sample representations from different methods for betterseparating disease and health control subjects, which is challenging if using single-cell transcriptomics directly. As shown in Figure 2 (d), which shows the sample embeddings, only sample embeddings from BrainBridge can separate heath control samples and AD samples in the embedding space. We considered quantitative metrics to evaluate the separation performance, including Normalized Mutual Information (NMI), Adjusted Rand Index (ARI), and Average Silhouette Width (ASW). These metrics are widely used in analyzing model performance for clustering [42, 43]. We average the scores of three metrics and report them in Figure 2 (e), which shows that embeddings from BrainBridge have the highest clustering scores. Therefore, BrainBridge can better represent disease state at the sample level.

### Performance Gain from training different phenotypes jointly

It is important to justify the contribution of training different phenotypes jointly is also, which not only allows us to use one model for various downstream applications but also allows learning across different predictions to potentially improve their performances. When developing a multi-task learning framework, we found that training age prediction and disease-state prediction together can boost the performance of each task, while training sex prediction with other phenotypes does not reduce performances. Therefore, it is useful to train disease-state prediction, age prediction, and sex prediction together to formulate a model with multiple targets, shown in Extended Data Figures 4 (a)-(c). Moreover, we explored the number of cells needed from each person for training. By treating disease-state prediction as the training target, we found that 500-1000 cells per subject can give us the best performance, whereas sampling more cells might lead to a performance drop, shown in Extended Data Figure 4 (d). This discovery aligns well with previous studies focusing on the bag size of multi-instance learning research [44]. Considering the noisy cells in scRNA-seq data, the choice of optimal number of sampled cells to gain the most useful information should be determined by experiments. Finally, considering the linkage between cell functions and disease effect, we also tested if only using cells from one cell type can help with disease-state prediction or not. Shown in Extended Data Figure 4 (e), among eight major cell types, using Interneuron (Inh) cells provides the highest prediction accuracy, while only using Microglia (Mic) cells provides the lowest prediction accuracy. However, prediction using individual cell types does not surpass 0.7 accuracy on the testing dataset, and such analysis cannot offer information to analyze cell-cell interaction. Therefore, we prefer multi-cellular modeling for making predictions and generating patient sample representations.

#### BrainBridge generates biologically meaningful representations of patients

Successfully integrating ROSMAP data and ABCA data can give us more insights about generating sample representations and prioritizing biological features. We visualize the cells from the ABCA data in Extended Data Figures 5 (a)-(d) with different labels, which convey similar information with the ROSMAP data: Raw transcriptomic profiles do not clearly separate AD subjects from health controls. We further utilize BrainBridge to generate patient sample embeddings and visualize the embeddings from the whole dataset jointly in Figures 3 (a)-(d) by different labels. Here we found that our sample embeddings are better at identifying biological signals, including disease states, age, and sex, while reducing the batch effect in the joint analysis of ROSMAP and ABCA.

**Fig. 3.**
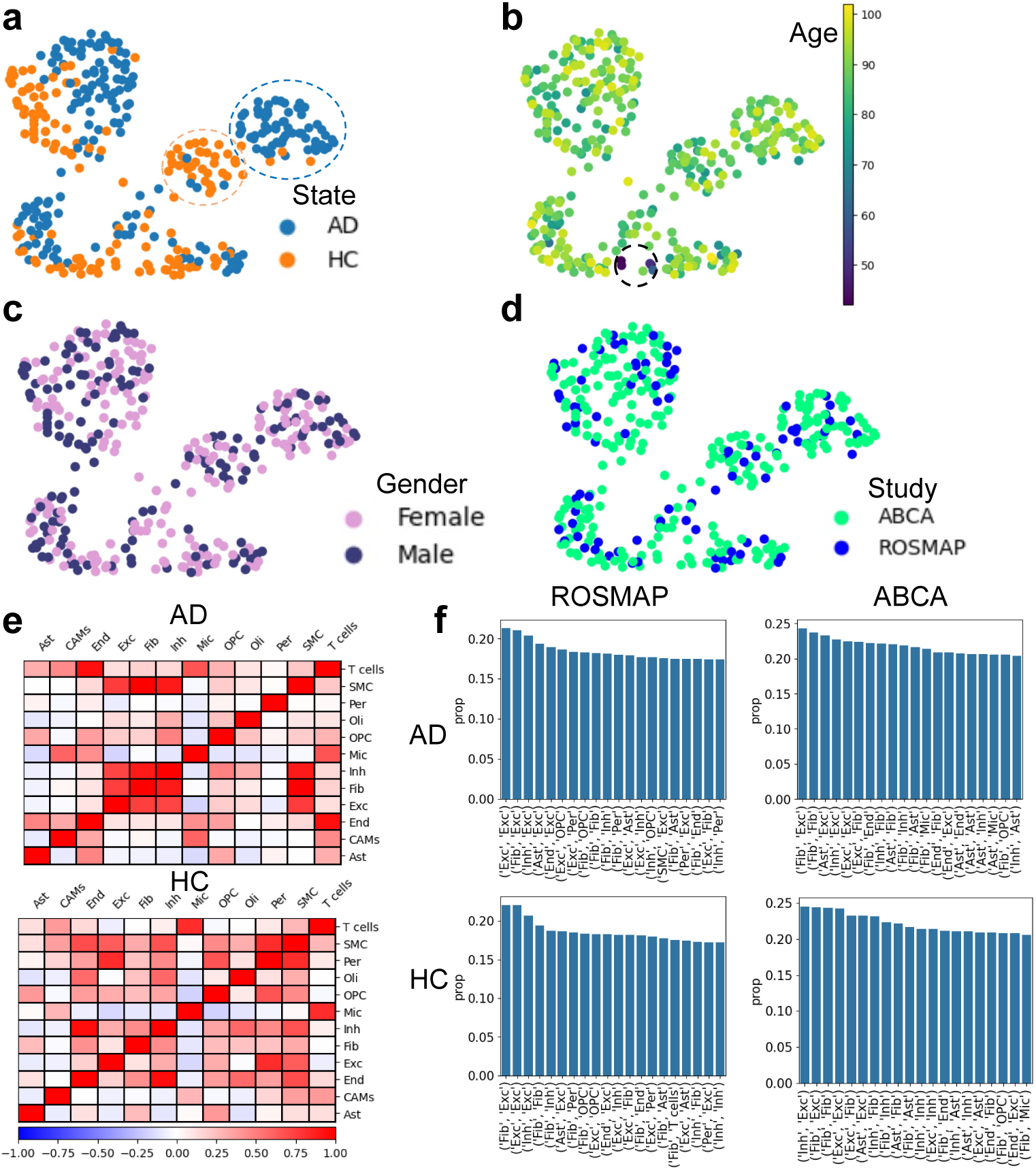
Analysis of patient sample representations and extracted cell-cell interactions. (a) UMAP plots of sample representations colored by disease states. We highlight disease-associated sample clusters and health-associated sample clusters with circles with corresponding colors. (b) UMAP plots of sample representations colored by age information. We highlight young-sample clusters with a black circle. (c) UMAP plots of sample representations colored by sex information. (d) UMAP plots of sample representations colored by sources of studies. (e) CCIs generated by the attention head of BrainBridge across samples from different states. (f) CCIs computed with CellPhoneDB based on single-cell transcriptomics from different studies for different disease states.

Therefore, BrainBridge can generate biologically meaningful sample embeddings, which allow us to detect isolated sample clusters to capture hidden information by the noise in the single-cell transcriptomic profiles, such as disease-specific signals. By focusing on the highlighted sample clusters in Figure 3 (a), we run the Leiden clustering algorithm [45] and computed the cell-cell interactions (CCIs) of expression profiles from these samples with the cellular attention from BrainBridge. The attention plots in Figure 3 (e) show that CCIs from samples with AD and HC present different patterns. CCIs among cell types including Inh, Fib, and Exc are obviously stronger in AD samples than in HC samples. To validate our discoveries, we also computed the CCIs based on the transcriptomic profiles with data from different studies to avoid batch effects as confounders, with results summarized in Figure 3 (f). We calculated the proportion of significant CCIs at the gene level between each cell-type pair using CellPhoneDB [46, 47] and presented the top 20 CCIs in the same figure. We found that Exc, Inh, and Fib also had high ranks compared with other CCIs. The enriched interactions between Exc cells and Inh cells in AD are also observed in other studies [48]. CellPhoneDB is supported by [49] as a reliable tool to estimate cell-cell interactions from scRNA-seq data. We observed consistent CCI patterns from BrainBridge and from CellPhoneDB in HC samples, and thus BrainBridge was able to capture the specific multi-cellular interactions in samples with different phenotypes, which further supports the idea of modeling patient samples by learning multi-cellular interactions. Moreover, BrainBridge can also work as an effective approach to extracting CCIs from atlas-level transcriptomic profiles.

### BrainBridge prioritizes biological features associated with diseases, aging effect, and sex effect

Similar to PaSCient, we utilized Integrated Gradients (IGs) [50] to compute the importance score of biological features’ contributions for prediction, but extending this design to fit multi-target prediction. By aggregating the importance matrix computed based on the transcriptomic of input samples, we can select important genes, cell types, and cells for interpreting the key factors affecting various phenotypes including disease states, aging effect, and sex effect.

We first extracted important cell types based on our importance scores for these three phenotypes in Extended Data Figure 6, which shows that Mic and T cells play an important role in biological activities related to different phenotypes, as they rank high in almost all plots of importance scores. Our discovery further highlights the importance of immune cells in responding to diseases as well as affecting the aging process [51–53]. Furthermore, Ast cells play an important role in AD samples, which aligns well with the studies of Ast cells’ functional change in AD patients [54]. Since the role of cell types is mainly determined by the gene programs, we further visualize the important genes of top three cell types associated with AD in Figure 4 (a). Top-ranked genes such as *FOXP2*, *GNLY*, and *SLC14A1* have been shown as network-driving genes in developmental disorders and neurodegeneration [55–57]. Moreover, these genes form gene programs positively associated with AD, and the relevance of these genes is further supported by identified Gene Ontology Enrichment Analysis (GOEA) pathways shown in Figure 4 (b) from Mic cells. We highlight the pathways specifically enriched in Mic cells and not in other cell types with biological mechanism interpretation using yellow stars, and notably, pathways such as regulation of microtubule depolymerization (GO:0031114) and positive regulation of microtubule polymerization or depolymerization (GO:0031112) are highly relevant, as microtubule stability and transport are critical in brain function and are known to be disrupted in pathology study of AD [58]. Cellular response to hexose stimulus (GO:0071331) and response to hexose (GO:0009746) suggest metabolic dysregulation, particularly in glucose metabolism, which is often observed in AD-affected brains [59]. Collectively, these enriched pathways suggest several hallmark features associated with AD.

**Fig. 4.**
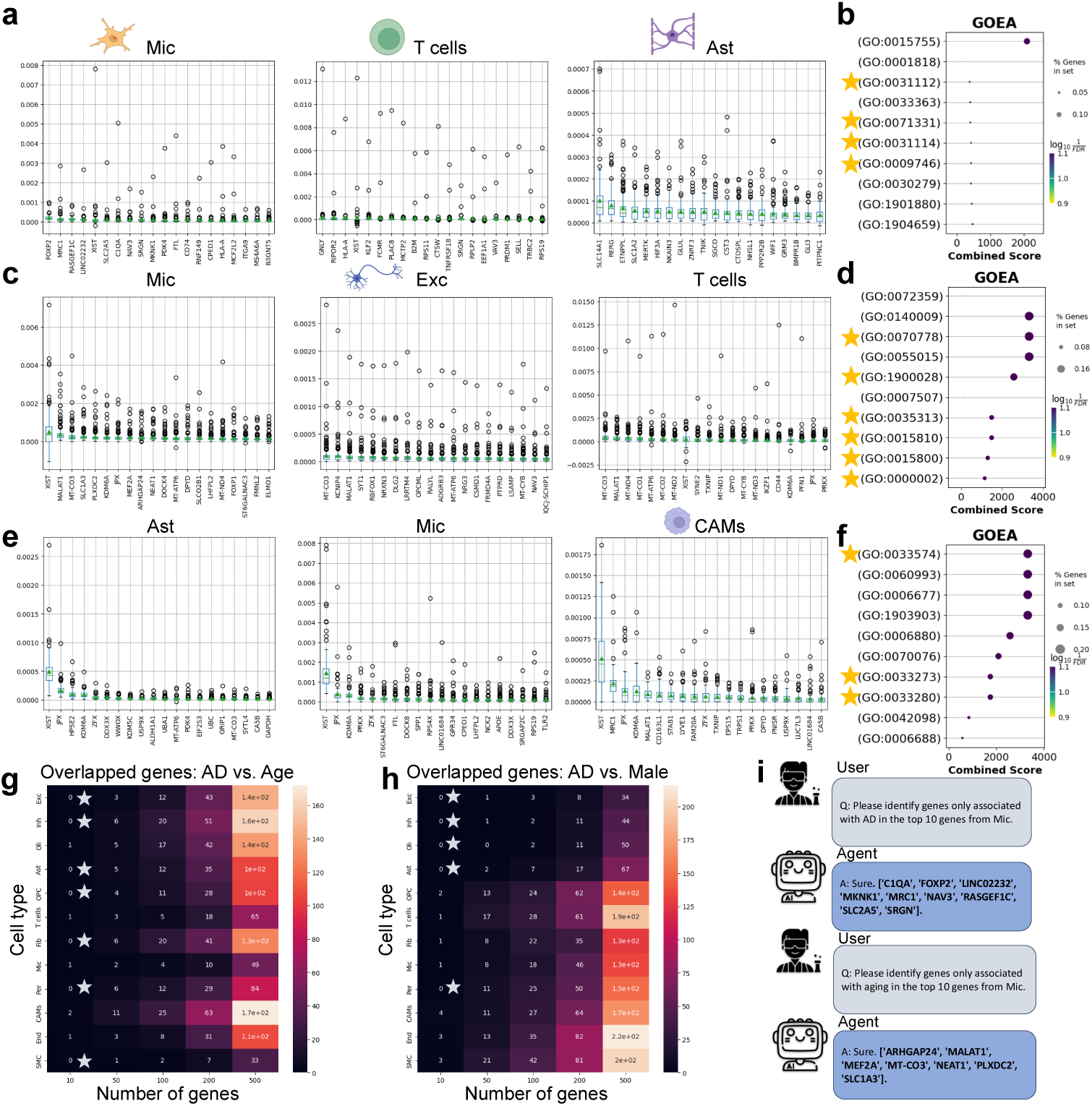
Interpreting gene programs associated with different phenotypes. (a) Important genes with importance scores associated with AD from the top three AD-associated cell types. (b) AD-associated GOEA pathways with gene set from Mic. (c) Important genes with importance scores associated with age from the top three age-associated cell types. (d) Age-associated GOEA pathways with gene set from Mic. (e) Important genes with importance scores associated with sex (male) from top three sex (male)-associated cell types. (f) sex (male)-associated GOEA pathways with gene sets from Mic. (g) Relationship between number of genes and overlapped genes between AD-associated genes and age-associated genes across different cell types. (h) Relationship between the number of genes and overlapped genes between AD-associated genes and sex (male)-associated genes across different cell types. (i) An example of using an agnetic AI system to extract genes specifically associated with AD by analyzing atlas-level transcriptomics. The yellow stars in (b), (d), and (f) represent validated pathways with literature support. The gray stars in (g) and (h) represent significant (p-value*<*0.05) cell types within the top 10 important genes with two-sided Fisher’s exact test. Cell-type logos are generated with biorender.com.

We also selected genes that might accelerate or decrease the speed of aging for the top three cell types, shown in Figure 4 (c). Based on our analysis, MT genes are identified in the top gene list for Mic, Exc, and T cells, which aligns with the role of MT genes in regulating the aging and programmed death process of cells [60]. We also plot the relationship between age and gene expression profiles for top-ranked genes of these three cell types in Extended Data Figure 7, which shows that the top-ranked genes generally have higher expression levels in the elder groups, and certain genes such as *XIST* (in Mic cells) and *MALAT1* (in T cells) have clear patterns of expression increasing with age increasing. MALAT1 is observed high expression in aging cells, which show that our analyses can capture reasonable patterns [61]. Figure 4 (d) shows the GOEA analysis of our gene programs associated with aging from Mic cells and yellow stars represent Mic-specific pathways. One key aging-associated process is mitochondrial genome maintenance (GO:0000002), which is critical for sustaining energy production and reducing oxidative stress—factors that are highly relevant to aging-related neurodegeneration [62]. Additionally, aspartate transmembrane transport (GO:0015810) and related amino acid transport processes such as L-aspartate transmembrane transport (GO:0070778) and acidic amino acid transport (GO:0015800) reflect metabolic adaptations that may shift with age to meet altered energy and biosynthetic demands [63]. The enrichment of wound healing, spreading of epidermal cells (GO:0035313) and negative regulation of ruffle assembly (GO:1900028) suggests roles for microglia in maintaining tissue integrity and modulating cell motility during inflammatory responses—functions that become increasingly dysregulated with aging [64, 65]. Overall, these enriched pathways underscore highly associated mechanisms that contribute to the biology of aging in the human brain.

We also identified sex-specific genes in the brain regions in the top three cells, visualized in Figure 4 (e). In our analysis, several genes located in the X chromosome, such as *XIST* and *JPX*, are identified. The expression of the *XIST* gene can inhibit gene expression from X chromosome [66], while *JPX* also works as a molecular switch in regulating gene activity in X chromosome [67]. Therefore, we can also model the difference of sex at the molecular level with BrainBridge. The selected pathways are summarized in Figure 4 (f) for Mic cells. Interestingly, the response to testosterone (GO:0033574) stands out as a key androgen-related pathway fundamental to male development, reproductive function, and secondary sexual characteristics [68]. The response to vitamin D (GO:0033280) and response to vitamin (GO:0033273) also play supportive roles, as vitamin D is known to influence testosterone levels and male reproductive health [69]. Therefore, these pathways collectively reflect biological signatures consistent with male-associated physiological and developmental status.

To identify genes that are only associated with targeted phenotype, we took the overlap of these genes as a shared functional gene group. For the whole gene programs that affect specific phenotypes, we subtracted genes from the shared group and generated disease-specific, aging-specific, and sex-specific gene programs. By alternating the number of selected genes to perform overlapping calculations, we record the possible change in Figure 4 (g) for AD vs. aging specific genes and Figure 4 (h) for AD vs. male-specific genes. These plots show that there are nearly no overlapped genes if we focus on the top 10 genes per cell type, and thus our identification of a specific gene set is practical and further supported by the non-significant result tested by two-sided Fisher’s exact test to examine overlapping. As the number of total selected genes increases, we also observe an increased number of overlapped genes, which is also reasonable. We also considered DESeq2 as a traditional baseline method. While DESeq2 can select two male-specific genes with adjusted p-value smaller than 0.05 without overlap with AD-specific genes, *AL358075.2* and *AC025265.2* are both long non-coding RNA without enough functional information. Traditional DEG method also cannot detect aging specific genes. Therefore, BrainBridge works as an alternative solution for detecting gene-phenotype associations for both discrete and continuous phenotypes.

In order to make it easier for users to extract the desired genes, we have implemented several functions to satisfy special gene requirements in the AI Agent system. Two examples of searching AD-specific genes and Aging-specific genes are summarized in Figure 4 (i), which clearly illustrates the convenience of using agent-powered BrainBridge.

### Refining information from other modalities with BrainBridge to perform multi-modal feature extraction

Results from Genome-Wide Association Studies (GWAS) [70] and Epigenome-Wide Association Studies (EWAS) [71, 72] have mapped genetic variants and epigenetic markers associated with human diseases/traits. These results are typically treated as different modalities, in comparison to the transcriptomic profiles considered in BrainBridge. Although GWAS and EWAS are conducted in cohorts with large sample sizes. The candidate regions tend to have many genes/sites and it is often challenging to pinpoint functional genes and could not directly provide risk factors associated with cell-type information. Therefore, leveraging the gene sets extracted from transcriptomic data may help refine the mapped genes from GWAS and EWAS with cell-type resolution and prioritize a smaller set of the most important genes. These genes are grounded by signals from additional data modalities and thus may have a stronger linkage with AD or the aging effect. The workflow pipeline of our method implemented in this section is summarized in Figures 5 (a) and (b).

**Fig. 5.**
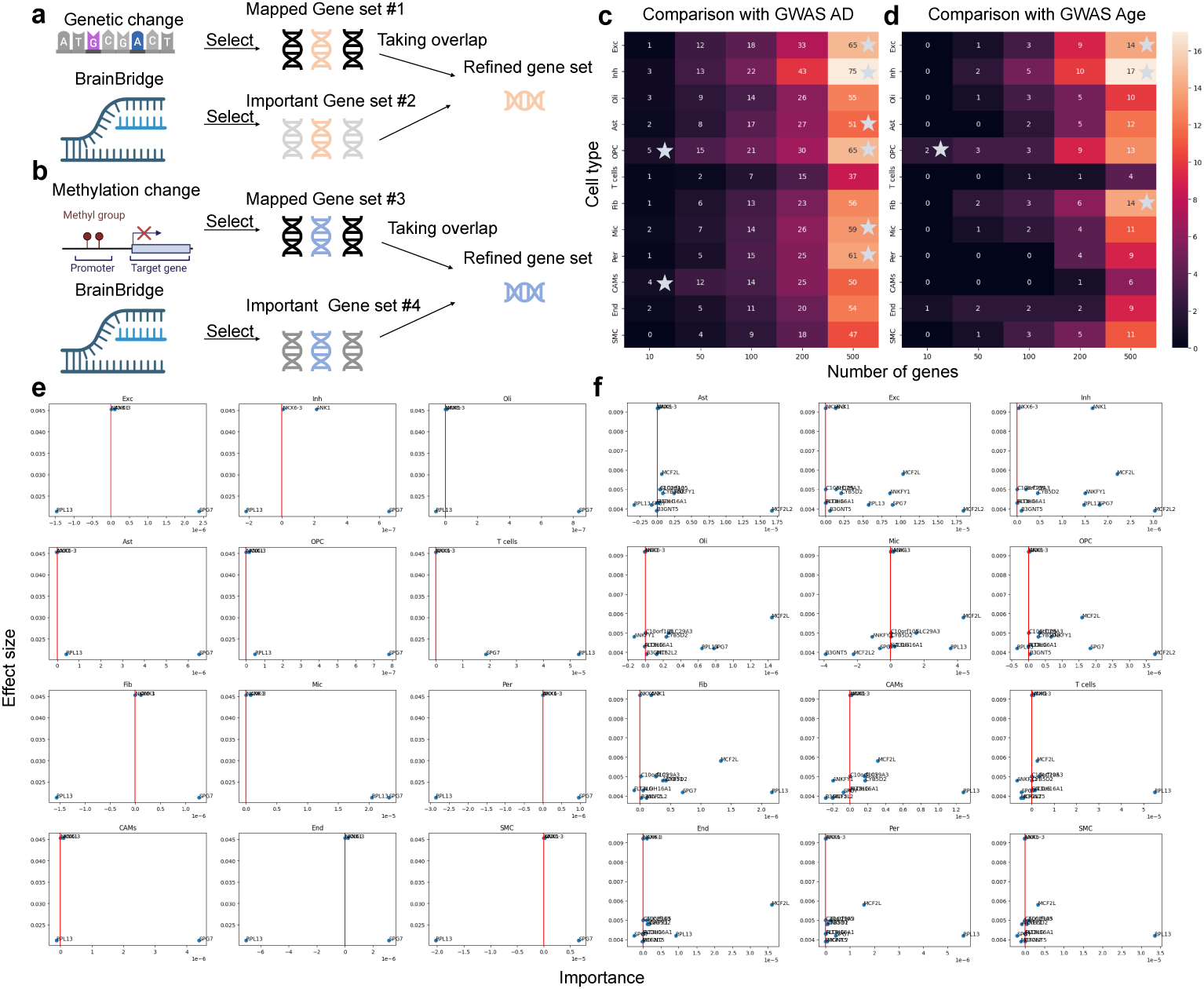
Gene set refinement with cell-type-specific resolutions. (a) Schematic workflow for refining gene sets from GWAS study with gene sets from BrainBridge. We took the overlap between two gene sets and generated the refined set. (b) Schematic workflow for refining gene sets from EWAS study with gene sets from BrainBridge. We took the overlap between two gene sets and generated the refined set. (c) Relationship between the number of genes and overlapped genes between AD-associated genes from GWAS study and BrainBridge across different cell types. (d) Relationship between the number of genes and overlapped genes between Aging-associated genes from GWAS study and BrainBridge across different cell types. (e) Relationship between effect size of AD-associated genes from EWAS study and important scores from BrainBridge across different cell types. (e) Relationship between effect size of Aging-associated genes from EWAS study and important scores from BrainBridge across different cell types. The gray stars in (c) and (d) represent significant (p-value*<*0.05) cell types within the top 10 and 500 important genes with the two-sided Fisher’s exact test.

We collected mapped gene sets associated with AD and the aging effect from the GWAS Catalog (1900 genes per set) [73]. According to Figure 5 (c), by analyzing the performance of refined gene sets in different cell types, we found that OPC cells and CAM cells demonstrated stronger enrichment in the top 10 genes. Moreover, when 500 genes were considered, OPC cells still had a strong enrichment, while Exc cells, Ast cells, Mic cells, and Per cells also showed significant overlap. Mic cells and Ast cells are also highlighted in our previous section. Exc cells and OPC cells also had the highest number of refined genes, which might raise further interest in analyzing the genetic change in these cell types for AD patients. OPC cells showed strong enrichment in the top 10 genes associated with the aging effect, shown in Figure 5 (d). Moreover, Exc cells, Inh cells, and Fib cells showed strong enrichment in the analysis of the top 500 genes, and thus we observed consistent enrichment enhancement by increasing the number of genes associated with AD or the aging effect. By analyzing the refined genes extracted from OPC cells with GOEA analysis shown in Extended Data Figure 8 (a), we identified pathways such as response to epinephrine (GO:0071871) and cellular response to epinephrine stimulus (GO:0071872), highlighting altered adrenergic signaling, which is known to affect neuroinflammation, glial activity, and cognitive processes in AD [74, 75]. Moreover, the enrichment of regulation of high voltage-gated calcium channel activity (GO:1901841) and regulation of actin filament-based movement (GO:1903115) point to disruptions in calcium homeostasis and cytoskeletal dynamics, both of which are well-established contributors to neurodegeneration [76, 77]. Furthermore, immune-related pathways such as engulfment of apoptotic cell (GO:0043652), apoptotic cell clearance (GO:0043277), and phagocytosis (GO:0006911) indicate active roles of oligodendrocyte progenitors in immune surveillance and clearance of neuronal debris, processes that are often impaired in Alzheimer’s pathology [78]. Extended Data Figure 8 (b) unravels the refined gene set associated with the aging process in OPC cells, and pathways such as postsynaptic membrane assembly (GO:0097104), postsynaptic density organization (GO:0097106), and presynaptic membrane organization (GO:0097090) reflect critical processes involved in maintaining synaptic architecture, which is widely acknowledged as the functional change in the aging process [79]. Similarly, neurotransmitter-gated ion channel clustering (GO:0072578) and positive regulation of neurotransmitter secretion (GO:0001956) indicate preserved or dysregulated synaptic signaling, which often deteriorates with age [80]. The negative regulation of dendritic spine development (GO:0061000) and neuron projection arborization (GO:0140058) suggest changes in neuronal morphology and connectivity, hallmarks of neural aging and cognitive decline [81]. Furthermore, the positive regulation of synaptic transmission, GABAergic (GO:0032230) also highlights the role of inhibitory signaling, which may become imbalanced in the aging brain [82]. Therefore, the refined gene sets provide biological signals for further exploring the negative effects brought by AD and the aging process.

We collected mapped gene sets associated with AD and aging effects from an EWAS research paper [83] with known effect size and significance. We only focused on the genes with positive effect sizes and p-values less than 0.05. To examine the coherence between important scores from BrainBridge and effect size from EWAS, we visualize the relationship between these two values with scatter plots across different cell types, shown in Figures 5 (e) and (f). Figure 5 (e) illustrates the heterogeneity of mapped genes across different cell types for analyzing the relationship between genes and AD. We identified *SPG7* as a hallmark in the refined gene set with consistent signs between effect size and importance score, which implicates *SPG7*’s important role in AD. Furthermore, *SPG7* has been shown to harbor both genetic and epigenetic risks associated with AD [84]. Another gene, *ANK1* is also supported by previous research as an AD-associated gene [85]. Figure 5 (f) shows the heterogeneity of mapped genes across different cell types for analyzing the relationship between genes and the aging process. We found a greater abundance of genes for aging compared to those associated with AD, and the refining results varied greatly across cell types. Here *RPL13* and *MCF2L* showed consistency between effect size and importance scores across most cell types, implying their roles in the aging process, which was supported by the literature [86, 87].

Therefore, analyzing atlas-level data can help us identify several factors associated with AD and the aging process, and refining gene sets can help us better understand and screen for specific types of genes. This may lead to more effective targeted drug development.

### Mapping prioritized genes towards 10X Visium data to capture the disease-associated spatial context

Considering the pathological changes induced by AD and aging, it is important to study the spatial contexts and expression levels of the genes identified through atlas-level data. Such external spatial transcriptomic data can also serve as an external validation of our discovery. Here we collected 32 human brain samples sequenced with 10x Visium technology [88] with different metadata to investigate the expression level validations of AD-associated genes and aging-associated genes under the spatial context with using spatial transcriptomics. We also utilized the cell-type deconvolution results of this dataset with SPOTLight [89] to perform spatial gene mapping in the cell-type resolution. Our pipeline is summarized in Figure 6 (a).

**Fig. 6.**
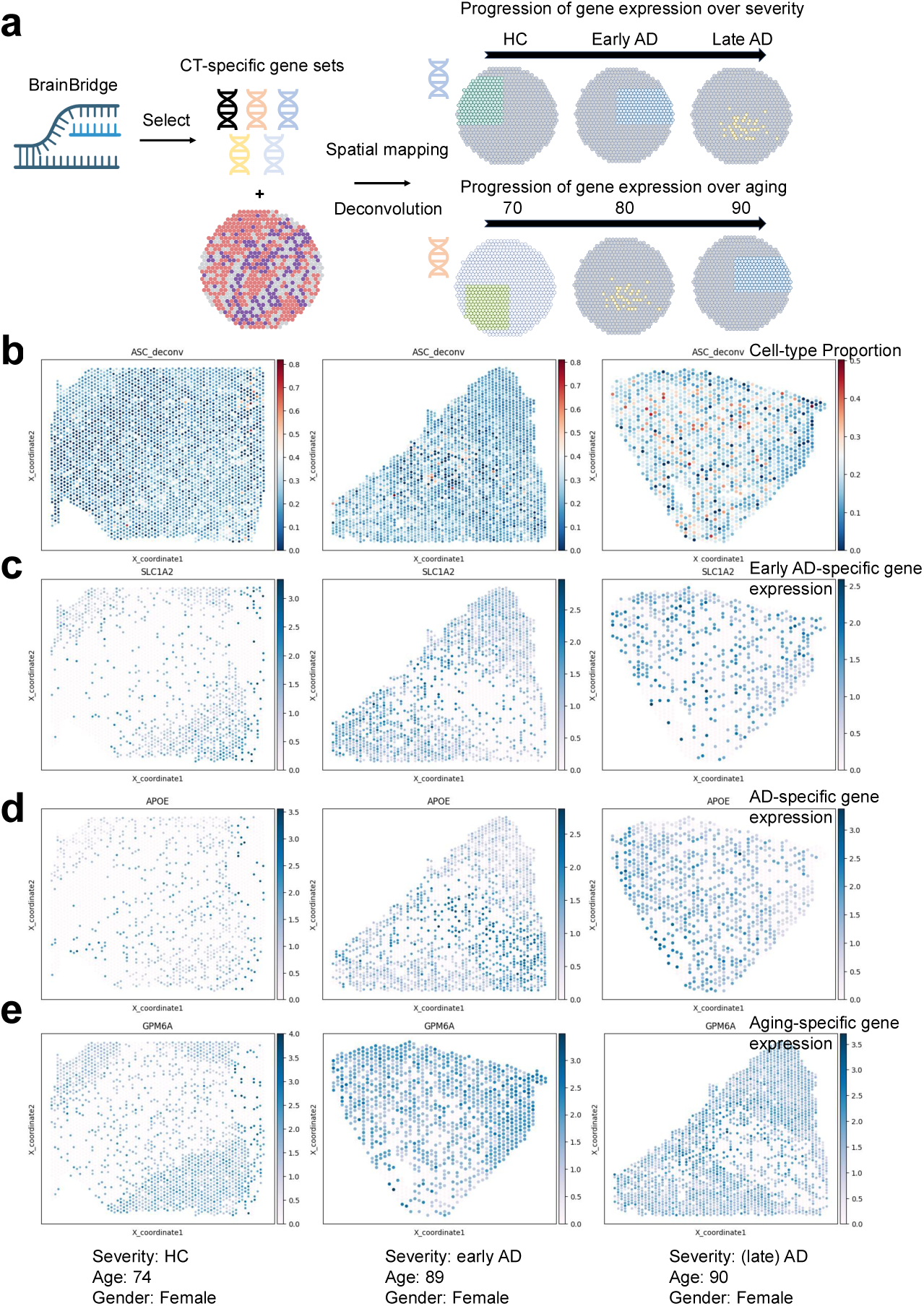
Mapping identified genes discovered from scRNA-seq to spatial transcriptomics. The information on age, sex, and health condition for the selected samples are recorded at the end of this figure. (a) Workflow pipelines of integrating BrainBridge with spatial transcriptomics for analyzing the expression profiles of AD- and aging-associated genes under spatial context. (b) Ast proportion under the spatial context after deconvolution based on the representative samples. (c) Expression profiles of *SLC1A2* under the spatial context based on the representative samples. (d) Expression profiles of *APOE* under the spatial context based on the representative samples. (d) Expression profiles of *GPM6A* under the spatial context based on the representative samples.

Here we focused on the associated genes from Ast cells and the corresponding deconvolution results are shown in Figure 6 (b). We observe that as the AD condition worsens, the centralized distribution of Ast in space becomes more pronounced. By examining the overlapped genes between highly expressed genes in early AD samples and AD-associated genes from BrainBridge, we identified *SLC1A2* as an early AD-associated biomarker. Figure 6 (c) shows the spatial expression levels of *SLC1A2*. The PCC between Ast proportion and *SLC1A2* expression levels was significantly positive, further supporting the consistency of Ast cells and *SLC1A2* expression levels. *SLC1A2* also had the most bright regions in the early AD sample. We also identified *APOE* as an AD-associated biomarker. Figure 6 (d) shows the spatial expression levels of *APOE*. The PCC between cell-type proportion and *APOE* expression levels is significantly positive, and the expression levels of *APOE* showed clear spatial patterns in the AD sample. *APOE* and *SLC1A2* were not highly expressed in the health control samples, which further supported the ability of BrainBridge in capturing correct AD-associated signals. Regarding the analysis of aging-associated genes, we identified *GPM6A* as a shared gene. The expression levels of this gene are shown in Figure 6 (e). We observed higher expression levels and a higher proportion of expressed spots in the elder groups. Therefore, the selected genes from BrainBridge are also validated in the spatial context as biomarkers for further analysis.

### Key contributions of BrainBridge for virtual cell analysis

In this manuscript, we trained a model with atlas-level single-cell transcriptomic profiles from the human brain and conducted several experiments to demonstrate the insights gained from our results. These contributions align with the call to build an AI-powered system for virtual cell analysis [90, 91]. To demonstrate this idea, we summarize the key discoveries by using BrainBridge as follows:

- BrainBridge works as an effective model in generating patient sample representations by encoding multi-cellular and phenotype-specific information, which can characterize biological signals such as disease-specific effects.
- BrainBridge prioritizes features including cells, genes, and cell types associated with different phenotypes, and further allows us to select specific gene programs.
- BrainBridge can integrate multi-modal information to interpret disease effect or sex effect, and the discovery is also validated under the spatial context.
- Powered by the Agnetic AI system, BrainBridge supports communication and interaction between researchers and models with natural language.

## Discussion

Given the growing scale of single-cell transcriptomic cohorts, it is crucial to develop methods that harness large sample sizes for phenotype-specific insights. Here, we introduce BrainBridge, which is inherited from our previous work PaSCient but focuses on the human brain atlas specifically and incorporates multi-instance learning, multitask learning, and semi-supervised learning for modeling the effects of AD, aging, and sex jointly. BrainBridge can not only generate patient sample representations by aggregating cell representations, but also prioritize cells, genes, and cell types associated with selected phenotypes jointly and individually. BrainBridge can also be used with data sources from other modalities such as GWAS, EWAS, and spatial transcriptomics to produce more robust results with external validation. Finally, we also deploy BrainBridge with an AI-powered agentic framework which can improve accessibility of BrainBridge.

BrainBridge works as an effective tool in uncovering and simplifying the complexity of single-cell gene expression in the human brain. First, by calling BrainBridge with inference mode, we generated sample representations which are enriched with disease-specific signals or health-specific signals that were previously masked by the noise and batch effect in the raw transcriptomics data. Second, BrainBridge prioritized gene programs associated with different phenotypes jointly or specifically, which provided multi-scale results to interpret disease effect, aging effect, and sex effect. For example, we identified gene *FOXP2* mainly associated with AD and gene *MALAT1* mainly associated with aging effect in Mic cells, which might inspire cell-type-specific therapeutic plans. Third, BrainBridge can also refine the gene sets from GWAS studies and EWAS studies, which are strong priors from other modalities and can further strengthen our inference results. Our findings implied that cellular heterogeneity might affect the function of these genes. Last but not least, we also provided spatially-resolved explanation of our detected genes with BrainBridge based on external validation datasets. We observed strong correlations between AD-specific genes or aging-specific genes and cell-type deconvolution outcomes, and found a rise in gene enrichment with worsening disease or aging. We also observed such gene enrichment in the RNAScope experiments performed in-vitro. Therefore, BrainBridge builds a bridge in extracting important information from atlas-level data and validating them with experimental methods, which aligns well with the community consensus of Artificial Intelligence Virtual Cell modeling.

Despite the contributions of BrainBridge, we also had challenges of applying and integrating BrainBridge for more purposes. First, BrainBridge does not directly model the spatial structures and we may consider other model architectures if we intend to model large-scale spatial transcriptomic data. This limitation might also affect our analysis for studying GWAS/EWAS. Second, due to the limitations of privacy and safety, it is hard to collect patient information in a human subjects and further extend it into multi-disease modeling. In the future, we plan to address the previous challenges by extending BrainBridge as a more flexible framework.

We are approaching the age of building foundation models for generating patient-centralized representations and boosting precision medicine. BrainBridge might unlock deeper insights into individual variability by integrating multi-modal biomedical data at scale. By capturing the full complexity of patient medicine, it enables more accurate predictions, tailored treatments, and advancements in healthcare delivery.

## Methods

### Problem Statement

Here we consider a set of (single-cell) transcriptomics with *n* profiles from atlas-level studies, denoted as 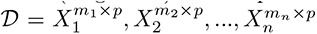, and the *k*th profile contains *m_k_* cells and *p* genes. Each profile corresponds to the recorded disease status *d_k_*, which is a binary variable. We also have profiles with chronological age information *t_k_*, which is a continuous variable; and sex information *g_k_*, which is also a binary variable. Our target is to train a model M, which accepts profiles from different samples as inputs and predicts the corresponding disease states as well as age information jointly with the help of representation learning, that is:

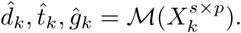

Here *s* represents the number of extracted cells from an individual sample.

### Model Architecture

The architecture contains three different components, including a cell-level encoder, a sample-level aggregator, and phenotype-specific decoders. We inherit the basic design from PaSCient [27], as the cell-level encoder is a Multilayer Perceptron (MLP)-based embedder and the phenotype-specific decoder is an MLP-based classifier. The sample-level aggregator is implemented with linear attention, that is, the attention weight represents the cell-level importance and we generate sample-level embeddings with a weighted-average method.

### Learning Disease Specificity and Aging Effect

When constructing the loss functions we use for model training, we utilize the cross entropy loss (CEL, L*_d_* and L*_g_*) to minimize the difference between predicted disease states and observed disease states as well as the difference between predicted sex and observed sex. We further use the mean squared error (MSE, L*_a_*) to minimize the difference between predicted chronological age and observed chronological age. Furthermore, we find that samples from the ROSMAP study [19] and the ABCA study [20] have different age scales, and thus we introduce a consistency regularizer (negative cosine similarity, L*_c_*) used in semi-supervised learning to ensure that the predicted age is consistent under different sampled cells from the same patient. For different loss functions, we have loss-specific pre-defined weights as *λ_d_*, *λ_a_*, *λ_g_*, and *λ_c_*, which correspond to the first three different loss function components. Therefore, our final loss function L*_f_* is:

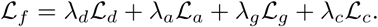

The weights are determined by observing the scales of specific loss functions to make a balanced optimization. We also compute the disease-state imbalance ratio of the training data and perform random oversampling [92] as a data augmentation approach at the sample level to make the disease states with a balanced distribution. In our implementation, *λ_d_* = *λ_a_* = *λ_c_* = 1, as their losses have similar scales. *λ_g_* = 0.0001 as age is a continuous variable and ranges from 20 to 102. We also investigate the contributions of hyper-parameter tuning as well as the introduction of matching loss L*_c_*, shown in Extended Data Figures 9 (a)-(c). We find that setting a learning rate around 1e-4 and a large batch size could improve model performance, and including matching loss can reduce the MSE between predicted age and observed age.

To reduce the cost of memory usage, we implement a new framework with a memory mapping design that can read large-scale transcriptomic data efficiently. By using this approach, we do not need to load the full matrix with metadata information to train the model, but only load the sampled cell by gene matrices and paired phenotypes for training/validation/testing.

### Generation of sample-level representations

When training BrainBridge, we leverage the idea of multi-instance learning (MIL) and treat the cells from one individual sample as instances, and these cells share the same phenotype information. To generate sample-level representations, we take the extracted cells *X_e_* and the encoder part of BrainBridge M*_e_*(), and thus sample representations of this individual can be computed with:

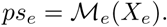

### Deep Learning Explainability for Feature Prioritization

To explain the predictions of our method, we implement the explainability with the integrated gradients (IG) method [50]. Each sample is represented as a matrix *X_k_* with shape (*m_k_,p*) and corresponding disease-state label *d_k_* (or age *t_k_* and sex *g_k_*). We also define a baseline sample 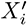 with the same shape but all values in 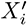 are 0. We write *F*(·) for the function mapping the sample matrix to the vector of predicted disease logits, and *F_j_*(·) for the *j*-th element of that vector, corresponding to the logit of the *j*-th disease/age information. In a binary classifier, *j* = 0 would correspond to the logit for control (i.e., health) prediction, and *j* = 1 to the logit for disease prediction. In a regressor, *j* is a constant. The output of IG applied to *X_k_* with baseline 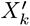 on the *j*-th dimension is given by an *attribution* matrix 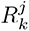 of the same dimensions as the input matrix. Each element of 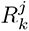 is defined as:

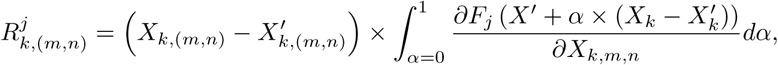

where 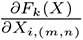 is the gradient of *F^k^*(·) along the (*m,n*) dimension, and *α* is used to generate the linear interpolation path between *X_i_* and 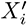. In practice, we use numerical integration to compute this quantity. Because the interpolation paths are identical for all dimensions (*m,n*), the importance can be computed with simple backward passes through the model.

By aggregating the importance matrix *R*, we can assess the contributions of cells, cell types, and genes for making predictions. Considering the sample size we can use to draw the conclusions, we focus on interpreting the contribution of cell types and genes in one specific cell type in this study. To interpret cell types, we record the cell-type information of instances in the given matrix and average the elements of the importance matrix by genes and cell types. To interpret cell-type-specific genes, we extract the cells with the same cell type from different samples, and average the elements of importance matrix by cells and samples.

For a specific cell type *c*, we can extract genes with positive attributions for AD denoted as 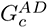, aging effect denoted as 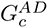, and sex effect denoted as 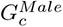 and 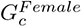. To compute shared and unique gene sets, we compute the overlap and difference of these gene sets. Therefore, we have:

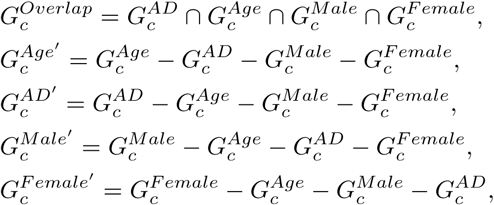

where 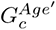 represents the gene set specifically associated with aging effect and 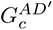 represents the gene set specifically associated with AD.

### Refining GWAS and EWAS

To refine the biomarkers identified with GWAS 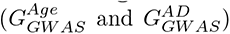 or EWAS 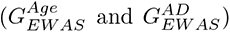 based on the results from BrainBridge, we first extract the effect size *β* estimated from the association study (if the study has such information), and then identify the gene sets with BrainBridge with the same sign of importance attribution. We then take the overlap of these two gene sets and define the newly generated gene sets as refined gene sets. We have analyzed the similarities and differences of gene sets from different studies in the Results section. Our approach can link transcriptomics with genetics and epigenetics for better interpretation.

### Mapping Prioritized Genes into Spatial Context

To validate the identified genes with external datasets with spatial context, we extract the cell-type-specific gene set and collect spatial transcriptomic datasets sequenced with 10X Visium technology [88]. The collected data contain three different disease states, including health control (HC), early AD, and AD. We utilize deconvolution results of SPOTLight [89] to identify cell-type-specific proportions of each spot, and further evaluate the consistency between extracted gene expression profiles and the distribution of cell-type proportions.

### Designing AI Agents for deploying BrainBridge

In order to better deploy BrainBridge and reduce the knowledge background required to use it, we design an AI agent with the OpenAI Agent framework, which allows users to easily prioritize features and generate sample embeddings. We define several tools, including a gene-searching tool, a cell-type-searching tool, and an embedding-generation tool. Our agent can call these tools based on input query prompts and further incorporate the output of the called function into the model-generated context. Moreover, implementing BrainBridge with an agent framework enables us to detect the potential problems in the questions and further suggests users with methods to correct the inputs.

**Details of RNA Scope study.** waiting… (it will be good to have RNAScope and perturb-seq results, if possible)

**Data preprocessing.** To preprocess transcriptomic data, we refer to the tutorials from Scanpy [35], filter out cells and genes with poor quality, and perform normalization followed by log-transformation for the raw count data to generate log-normalized data. To train the model, we did not select highly variable genes, which enable its potential to perform whole-genome modeling.

**Explanations of baseline models.** In this section, we introduce the baseline methods.

- MultiMIL [24]: MultiMIL is a model trained with weakly supervised design to predict the phenotype annotation of multi-omic datasets from different studies. It can return the predicted phenotype annotation after training based on the testing dataset and does not directly generate patient representations.
- Scimilarity [36]: Scimilarity is pre-trained with large-scale single-cell transcriptomic datasets to annotate cells and query cells with similar cell states. It can generate cell embeddings and these embeddings are later averaged into patient representations, by following the pipeline provided in PaSCient.
- MrVI: MrVI [25] is a deep generative model used to analyze sample-level heterogeneity, which may capture clinically relevant stratifications of patients. We average the cell embeddings learned from MrVI to generate patient representations.
- pseudobulk-exp: pseudobulk-exp means we formularize patient representations by averaging the gene expression profiles of different cells within the same sample directly. It is a simple baseline by following the pipeline provided in PaSCient.
- mean age: mean age represents the baseline used for benchmarking age prediction. We take the average age based on the testing dataset and compare it with the results from other methods.

**Explanations of metrics.** In this section, we introduce the information on metrics we used in our evaluation.

To compare the difference between predicted variable and observed variable, we utilize the Accuracy and Weighted F1 score for evaluation. Both scores are in [0,1] and higher scores mean better model performance.

To compare the difference between predicted age and observed age, we utilize Pearson Correlation Coefficients (PCCs) and Mean Squared Errors (MSEs) for evaluation. PCC is in [-1,1] and higher PCC means better model performance. If the predicted age is a constant for different samples, we reported -1 as the PCC. MSE is in (0,∞) and higher MSE means worse model performance.

To compare the gene sets extracted from BrainBridge and the gene sets covered by previous studies, we utilize Gene Ontology Enrichment Analysis (GOEA) and Fisher’s exact test. (Adjusted) P-value threshold of significance is 0.05.

## Dataset availability

All of the datasets used in this manuscript are publicly available. Accessing ROSMAP datasets requires an application. We summarize the dataset statistics and download information in Supplementary File 2.

## Code availability and reproductivity

We rely on Yale High-performance Computing Center (YCRC) and utilize one NVIDIA H100 GPU with up to 150 GB RAM for training and inference.

The codes of BrainBridge can be found in https://github.com/HelloWorldLTY/brainbridge. The license is MIT license.

## Acknowledgments.

We thank Dr. Jia Zhao for her suggestions to improve the quality of our manuscript.

## 5 Author contributions

waiting…

## 6 Competing interests

The authors declare no competing interests.

---

**Extended Data Fig. 1.**
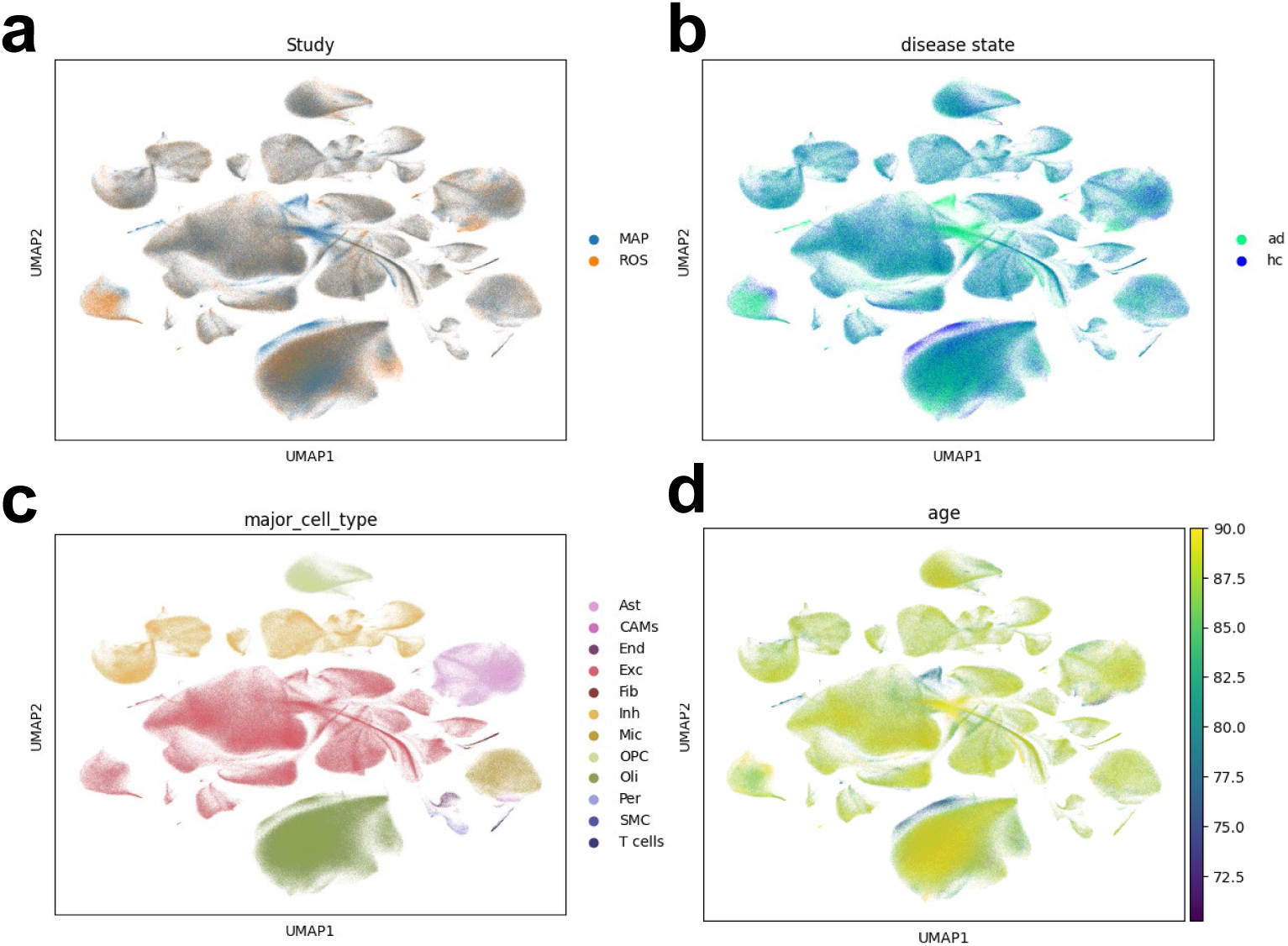
_Visualization of cellular information from ROSMAP. (a) UMAP plots of cells colored by sources of study (ROS or MAP). (b) UMAP plots of cells colored by disease states. (c) UMAP plots of cells colored by cell types. (d) UMAP plots of cells colored by age information._

**Extended Data Fig. 2.**
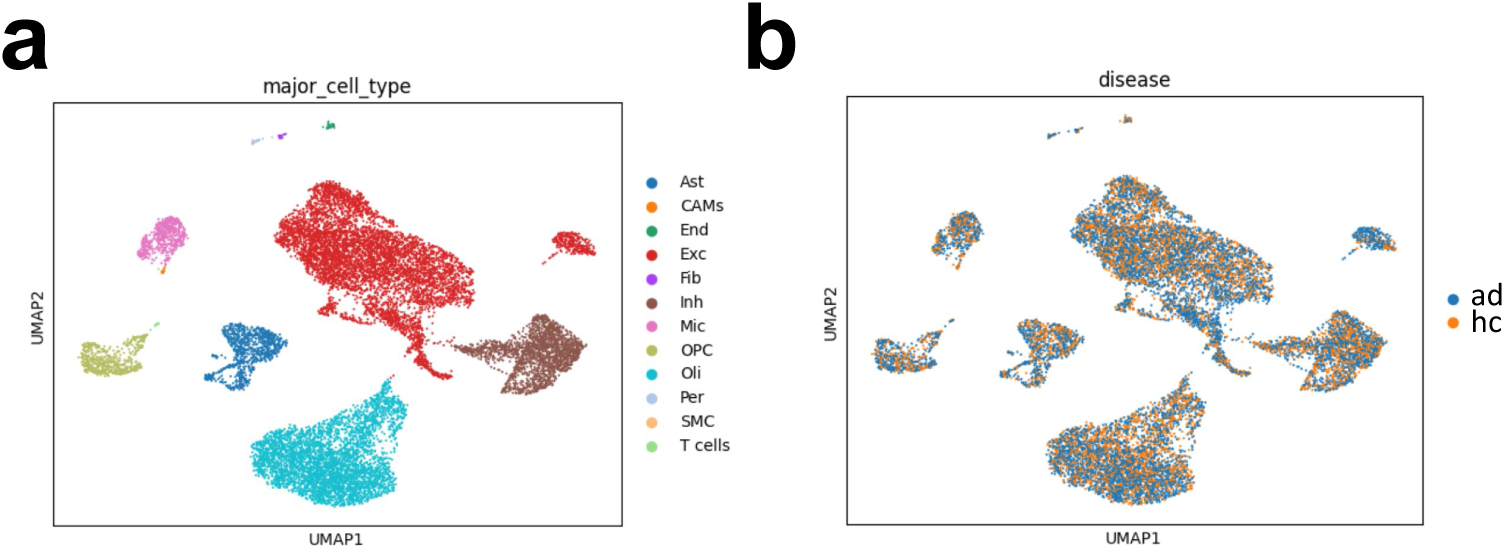
Visualization of cell embeddings generated by Scimilarity from ROSMAP. We subsampled datasets into 10% for more efficient processing. (a) UMAP plots of cells colored by cell types. (b) UMAP plots of cells colored by disease states.

**Extended Data Fig. 3.**
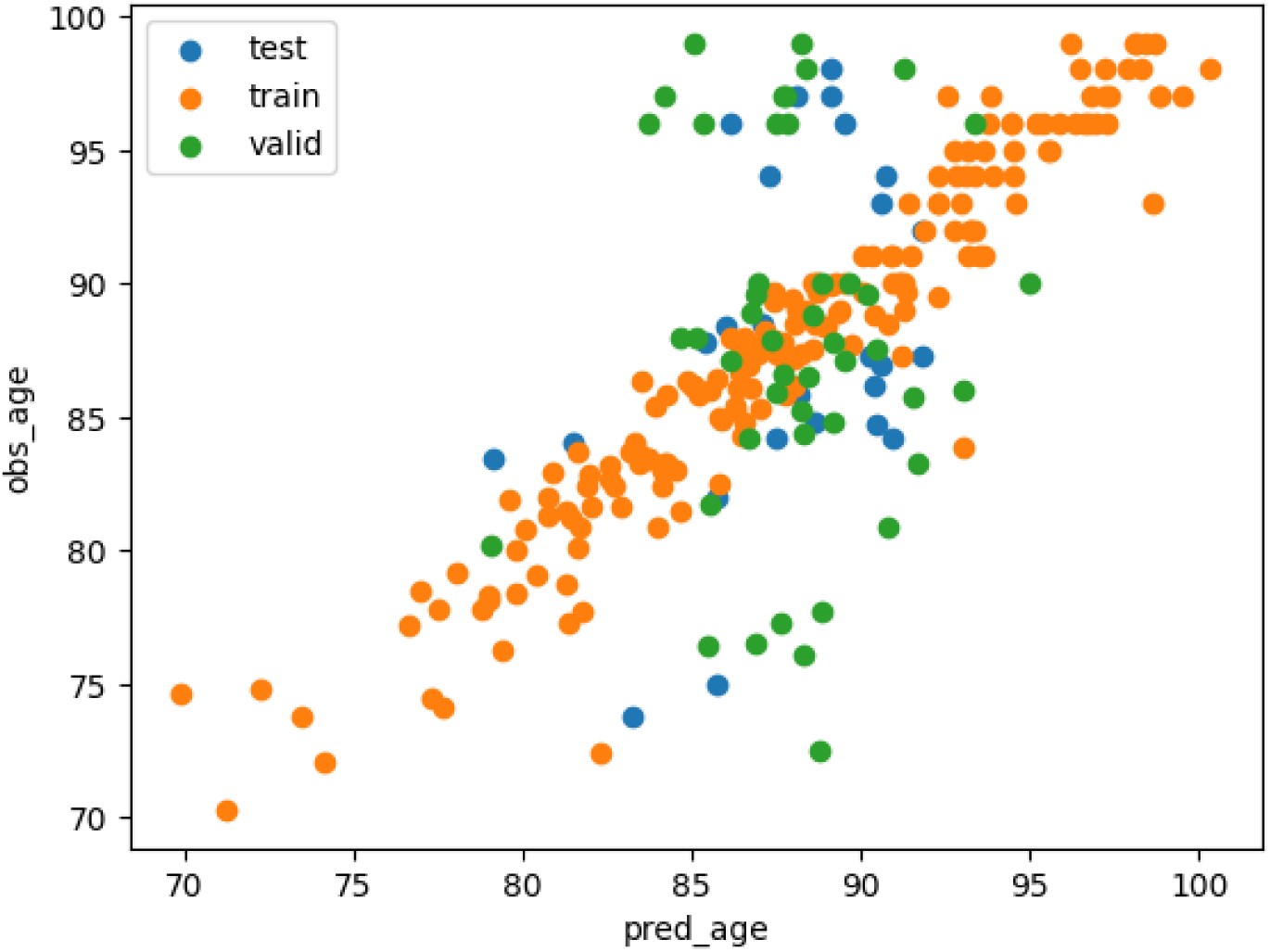
Scatter plot to visualize the relationship between predicted age and observed age for ROSMAP data.

**Extended Data Fig. 4.**
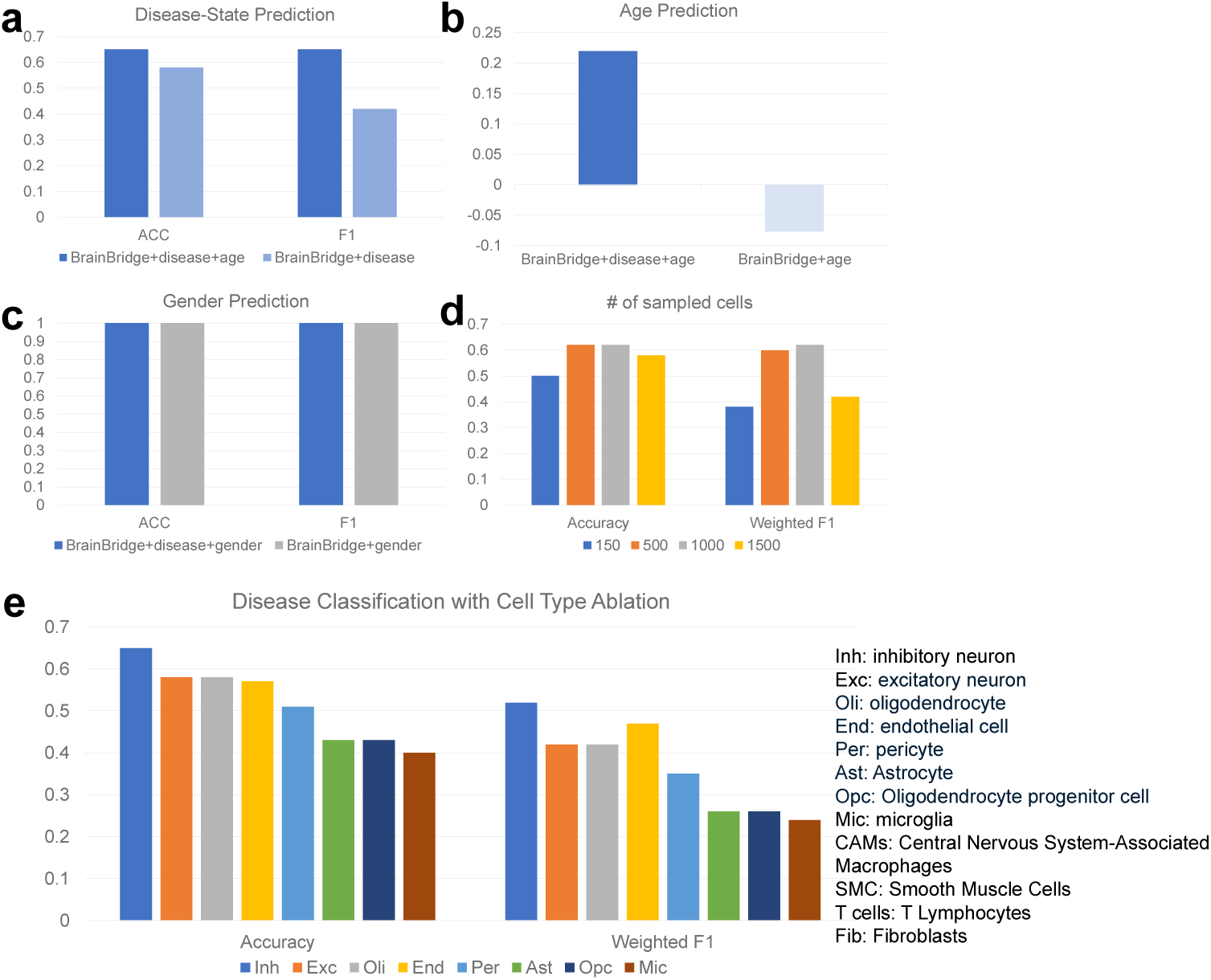
Ablation studies of BrainBridge. (a) Results of disease-state classification to compare the joint prediction mode and the disease-only mode. (b) Results of age regression to compare the joint prediction mode and age-only mode. (c) Results of sex classification to compare the joint prediction mode and the sex-only mode. (d) Results of disease-state classification under different number of cells sampled to formulate the patient representation in the training stage. (e) Results of disease-state classification by only using one cell type.

**Extended Data Fig. 5.**
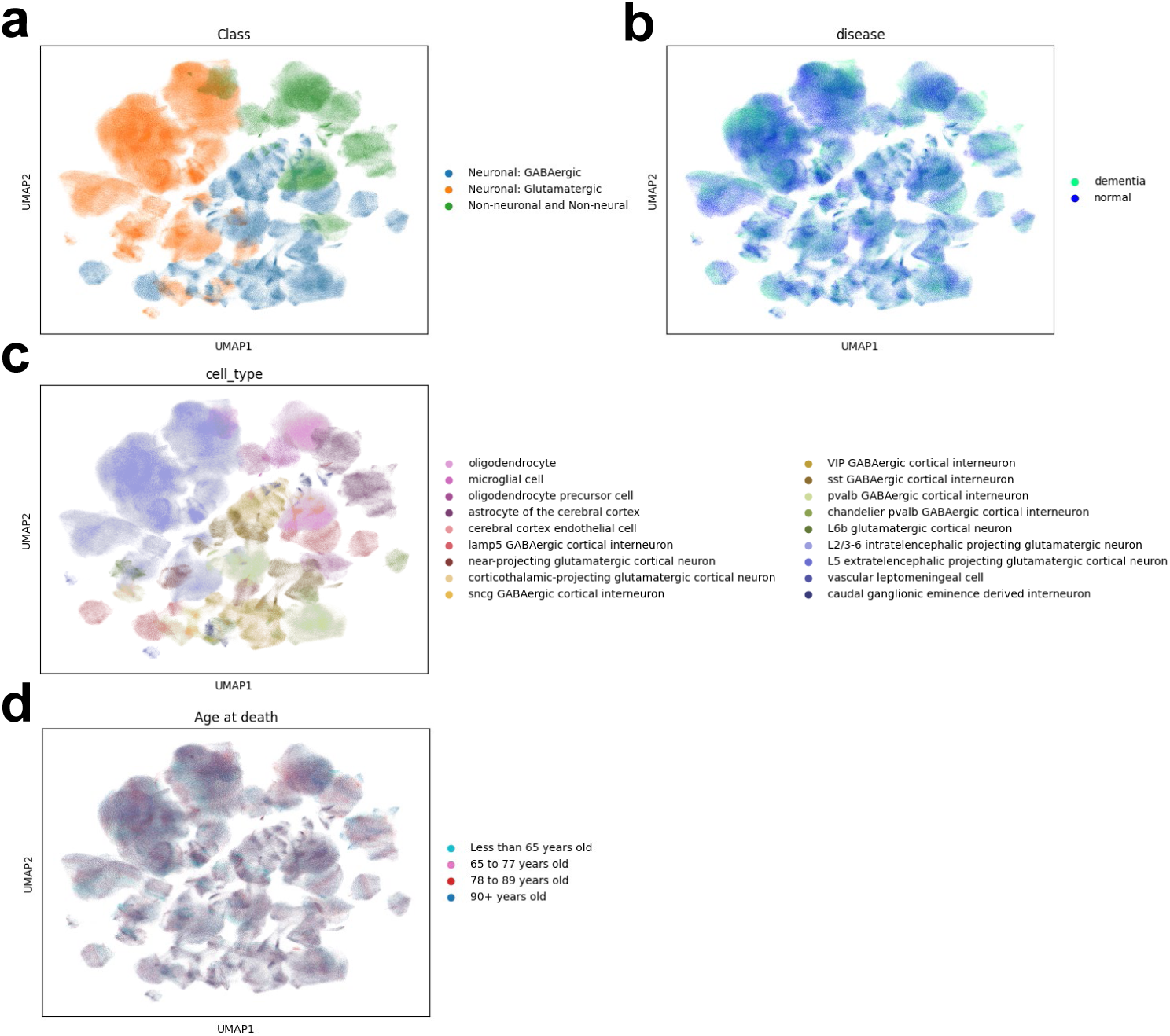
Visualization of cellular information from ABCA. (a) UMAP plots of cells colored by general cell types. (b) UMAP plots of cells colored by disease states. (c) UMAP plots of cells colored by cell types. (d) UMAP plots of cells colored by age information.

**Extended Data Fig. 6.**
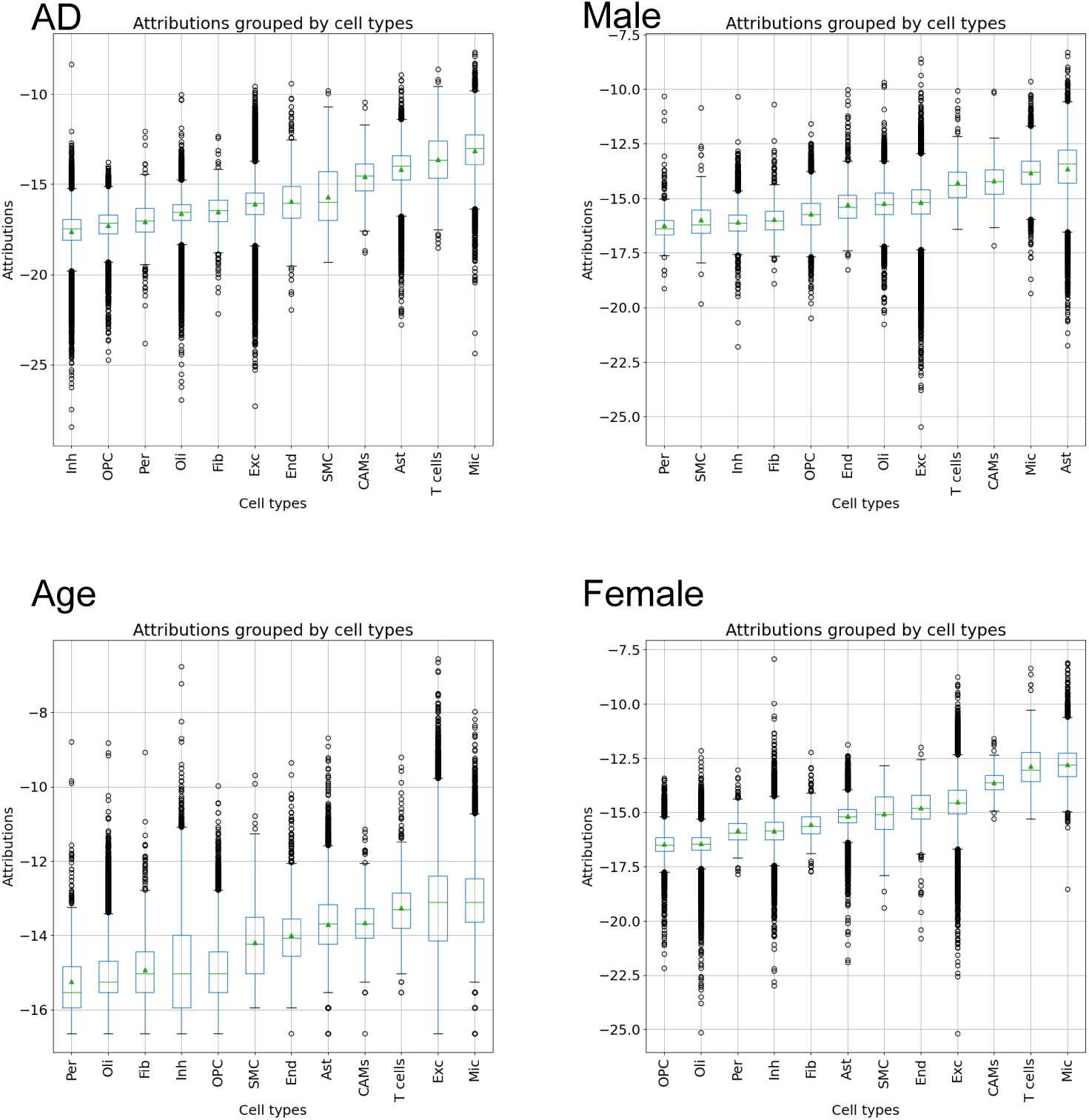
Important scores across different cell types targeting on different phenotypes, including AD, Aging, Male, and Female. The scores are performed after log-transformation.

**Extended Data Fig. 7.**
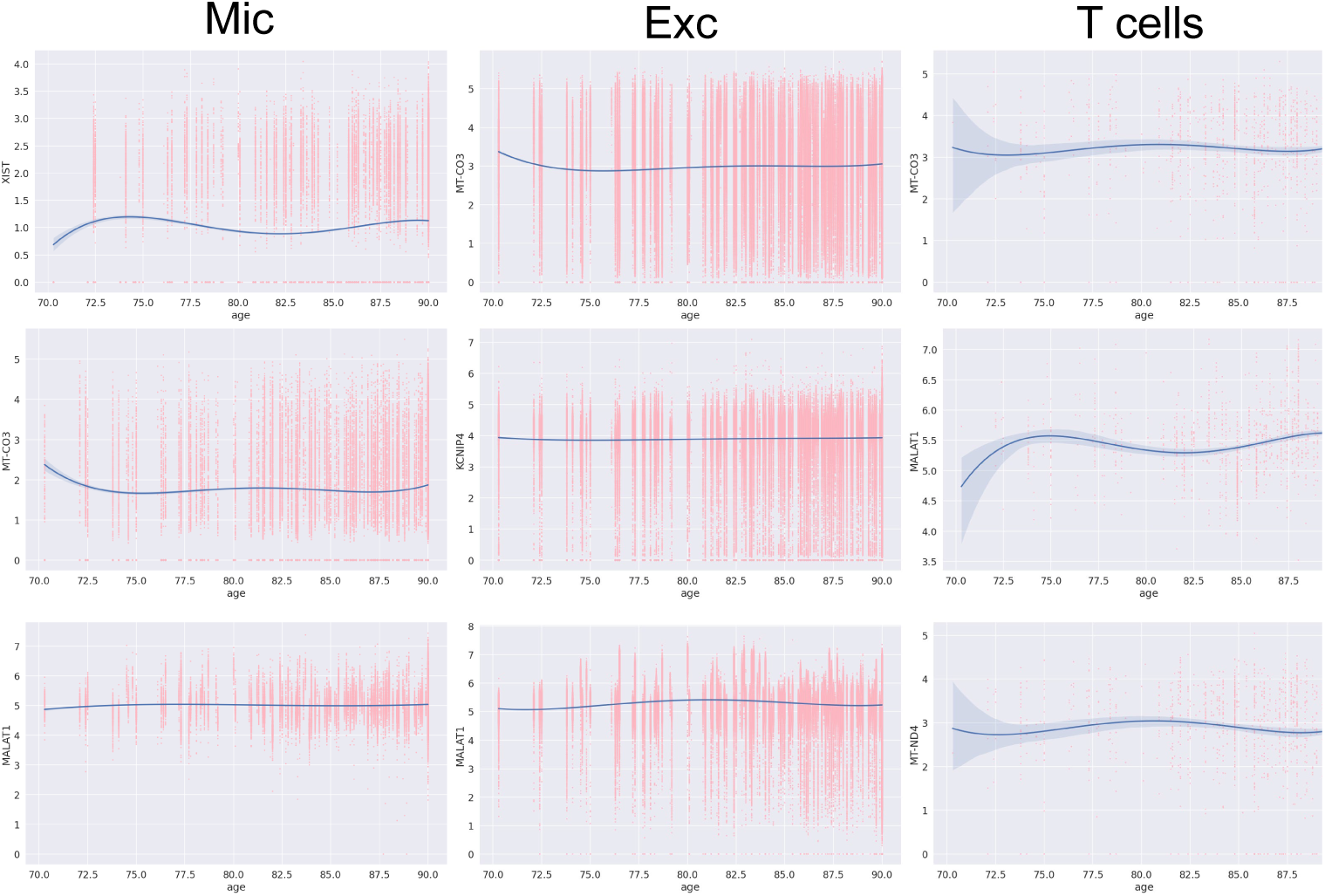
The relationship between gene expression profiles and aging process for top-ranked gene sets across top-ranked cell types associated with aging effect.

**Extended Data Fig. 8.**
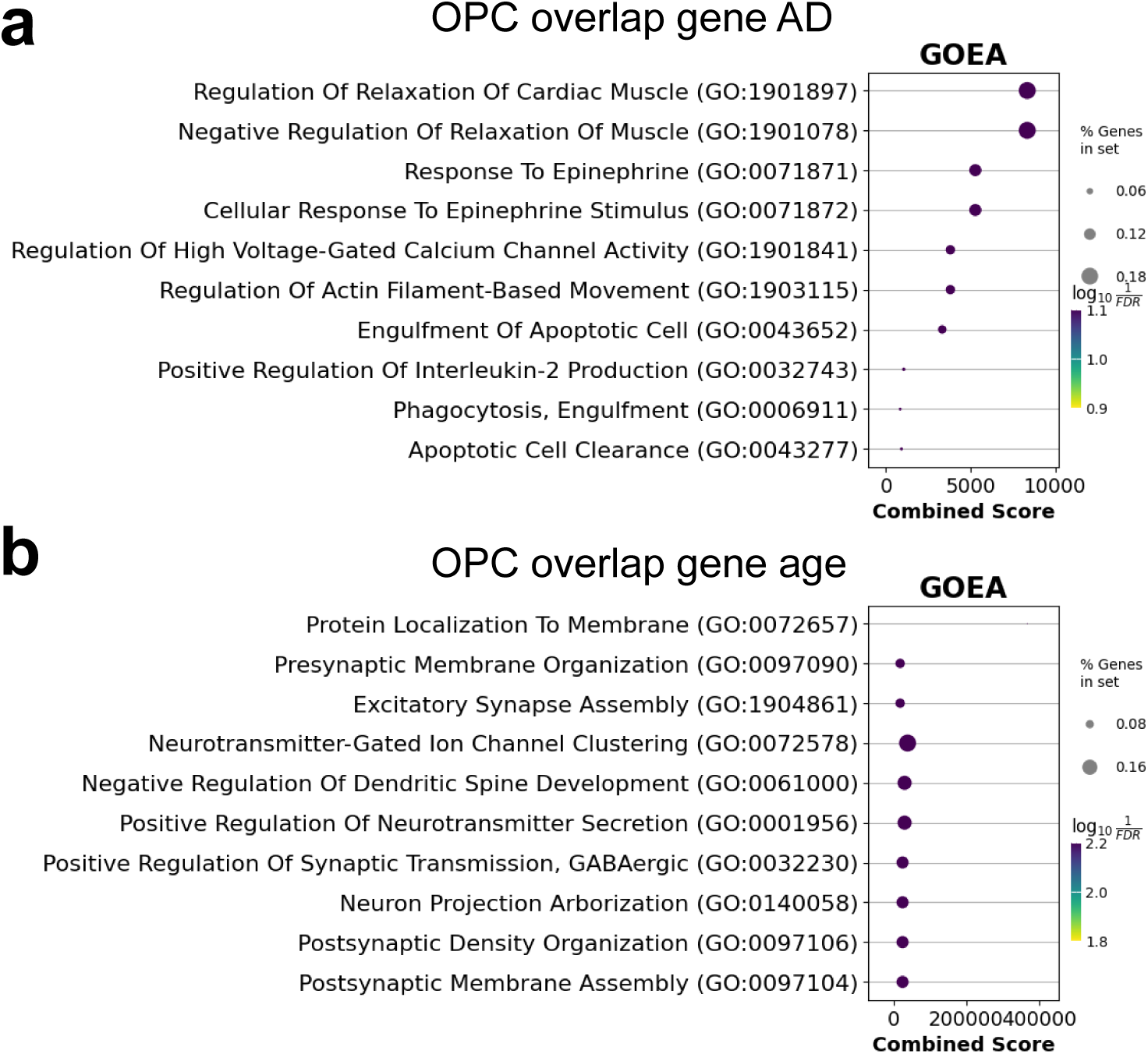
GOEA results after refinement with GWAS resources. (a) represents identified pathways from the OPC cells related to AD. (b)represents identified pathways from the OPC cells related to aging.

**Extended Data Fig. 9.**
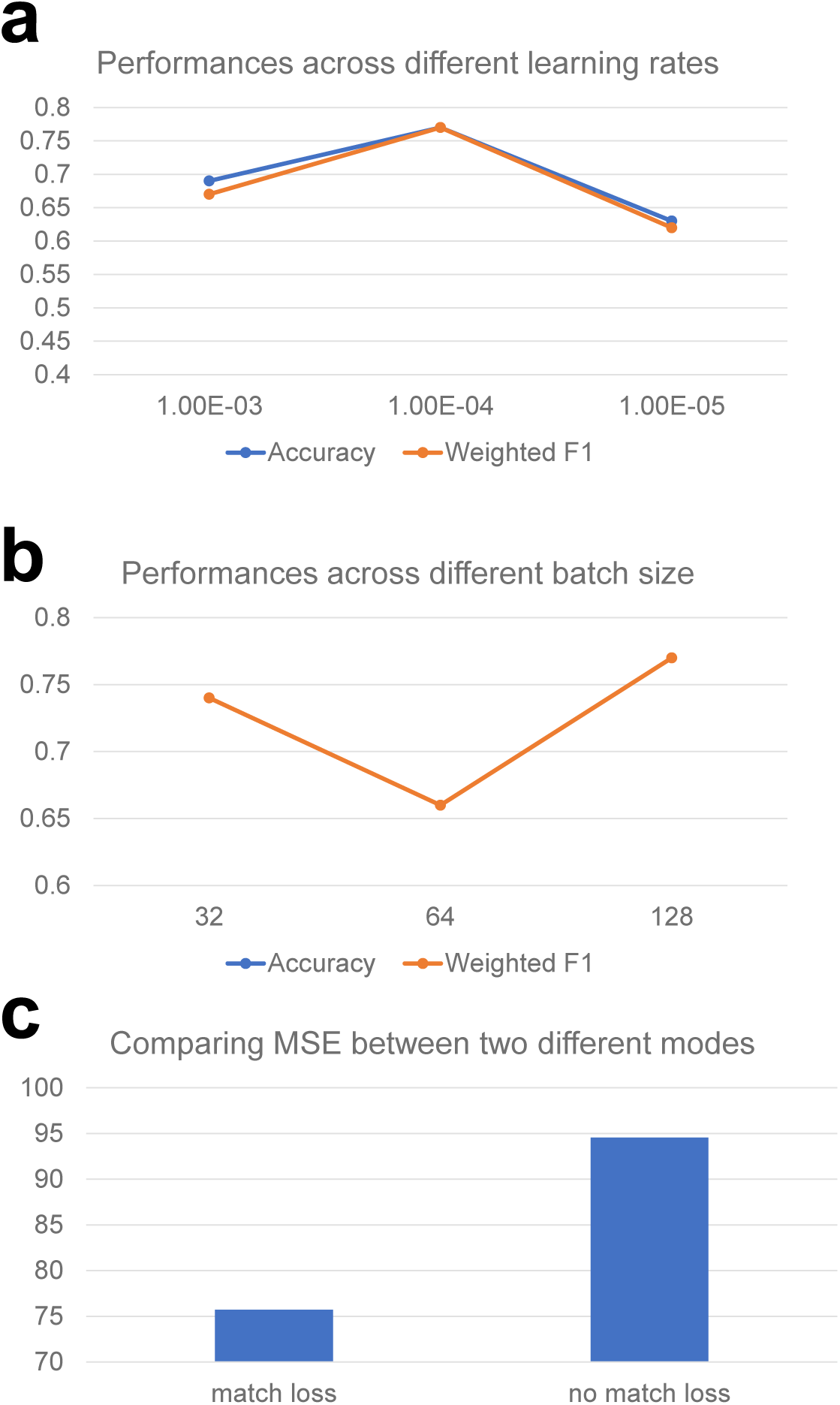
Results of hyper-parameter turning and ablation studies for the loss function. (a) Relationship between learning rate and disease-state prediction performance. (b) Relationship between batch size and disease-state prediction performance. (c) MSE of age prediction for two different modes of BrainBridge, as one has matching loss and the other one does not have matching loss.

## Notes

### Competing Interest Statement

The authors have declared no competing interest.

